# Organelle partitioning in the multi-budding yeast *Aureobasidium pullulans*

**DOI:** 10.64898/2026.04.17.719237

**Authors:** Alison C.E. Wirshing, Michelle Yan, Daniel J. Lew

**Affiliations:** Department of Biology, Massachusetts Institute of Technology, Cambridge, MA 02139

## Abstract

Cellular organelle content is fairly constant within a given cell type. This is accomplished in part by ensuring equitable organelle partitioning during division. Much of our understanding of organelle inheritance has come from investigating cells that divide in half producing two daughter cells. However, more elaborate division strategies that give rise to multiple daughters are not uncommon in nature. Here, we present the first characterization of organelle inheritance in a fungus that grows by multi-budding, producing several (2-20) daughter cells in a single cell cycle. We find that some organelles (mitochondria and ER) are evenly delivered to all growing buds, while others (vacuole and peroxisomes) are more variably inherited. We discuss the implications of even and uneven inheritance for this polyextremotolerant fungus capable of growing in dynamic, and diverse, environments.

## INTRODUCTION

Different cell types have quite distinct and characteristic organelle contents (Mostafa, 2022; Rhoads *et al*., 2025). However, for a proliferating population of a particular cell type, the cells appear to contain quite homogeneous amounts of each organelle, suggesting that there is an optimal organelle composition (Marshall, 2020). The functional consequences of perturbing organelle abundance have mainly been examined in extreme cases where no organelle is inherited (e.g., (Jin and Weisman, 2015)), although more modest changes in organelle abundance may also impact cell fitness (Chacko *et al*., 2025; Ray *et al*., 2025). Many symmetrically dividing cells adjust cell shape and organelle distribution so as to minimize inequalities in organelle inheritance at cell division (Ramkumar and Baum, 2016; Champion *et al*., 2017; Glotzer, 2017; Carlton *et al*., 2020; Chacko *et al*., 2023; Kors and Schlaitz, 2024). Disrupted organelle inheritance can delay cell cycle progression, enabling correction of organelle content (Jin and Weisman, 2015; Leite *et al*., 2023; Chacko *et al*., 2025). In addition, heterogeneity in organelle composition is a frequent hallmark of disease (Chang and Marshall, 2017; Patat *et al*., 2024). These findings suggest that equal partitioning of organelles among daughter cells is important, although the specific reasons are often unclear.

Homogeneous organelle partitioning is harder to achieve for cells that divide asymmetrically, like budding yeasts. Nevertheless, studies on the model yeast *Saccharomyces cerevisiae* have found that for various organelles, organelle size (volume or surface area) is maintained as a relatively constant proportion of total cell size (Jorgensen *et al*., 2007; Uchida *et al*., 2011; Rafelski *et al*., 2012; Chan *et al*., 2016). During bud growth, yeast cells use polarized actin cables to deliver mitochondria (Westermann, 2014), ER (Du *et al*., 2004), Golgi (Rossanese *et al*., 2001), vacuoles (Weisman, 2006), and peroxisomes (Fagarasanu *et al*., 2010) to the bud. Each organelle uses specific adaptor proteins to attach the organelle to a type V myosin that walks along the actin cables (Knoblach and Rachubinski, 2015). How such cells determine how much of each organelle to deliver to the bud and how much to retain in the mother is largely unknown.

Unlike *S. cerevisiae*, some budding yeasts can make more than one bud within a cell cycle (Restrepo and Jiménez, 1980; Mims *et al*., 1988; Lubbehusen *et al*., 2003; Mitchison-Field *et al*., 2019; Henß *et al*., 2022). Making several buds at once raises the question of how fairly the mother cell’s contents are apportioned among the sibling daughters. If a specific organelle composition is important for daughter cell function, one could imagine checkpoint-style pathways that enforce equal inheritance. Alternatively, cells may thrive with a wide variety of organelle compositions, in which case considerable variability might be allowed. Because organelle composition adapts to differing nutrient conditions (Sakai *et al*., 2006; Van Der Klei *et al*., 2006; Di Bartolomeo *et al*., 2020), it is also possible that distinct organelle compositions might pre-dispose daughter cells to thrive in different environments.

*Aureobasidium pullulans* is a polyextremotolerant multi-budding yeast that thrives in a broad range of environments (Gostinčar *et al*., 2011). For a generalist species in changing environments, daughter cell heterogeneity might represent an adaptive form of bet-hedging (Bagamery *et al*., 2020). This led us to investigate the degree of heterogeneity in organelle inheritance among the daughters of this multi-budding yeast.

In recent work, we addressed how *A. pullulans* mother cells apportion growth (Wirshing *et al*., 2025) and nuclei (Wirshing *et al*., 2024; Petrucco *et al*., 2025) among sibling daughters. Our findings suggested that equalization of polarity factors among the nascent bud sites led to development of similar actin networks directed towards each bud, leading to the growth of daughter cells of similar size. Moreover, nuclear distribution during mitosis usually resulted in each daughter receiving a single nucleus, with the remaining nuclei staying in the mother. However, there were quite frequent exceptions to these trends. About 5% of mothers made buds that differed in volume by >50% (Wirshing *et al*., 2025). And in ∼10% of cases, sibling buds received unequal numbers of nuclei (e.g., two nuclei versus one) even though daughter cell volumes were similar (Wirshing *et al*., 2024; Petrucco *et al*., 2025). Thus, sibling cells can be born with somewhat different size and/or nuclear composition.

Here, we examine how various organelles are partitioned across multiple sibling buds in *A. pullulans*, and uncover the natural degree of cell-to-cell variation in organelle composition among isogenic populations of proliferating cells of an organism that exhibits high morphological plasticity. This enables comparisons with organelle composition variability in *S. cerevisiae* (Rafelski *et al*., 2012; Mukherji and O’Shea, 2014; Chan *et al*., 2016) and sets the stage for a comparative analysis of mechanisms that promote homogeneity and/or variability in organelle composition.

## RESULTS

### Morphology and dynamic behavior of organelles in *A. pullulans*

To characterize organelle inheritance in *A. pullulans* we introduced fluorescent tags at endogenous loci of genes predicted to encode proteins localizing to various organelles. Cytosolic 3xmCherry in the same strain, described previously (Wirshing *et al*., 2025), enabled cell segmentation and analysis of organelle composition as a fraction of total bud or cell volume. Mitochondria were visualized using the previously validated probe, Cit1-GFP (Wirshing *et al*., 2024). To visualize other organelles, we tested 2-4 resident proteins each for the ER, vacuole, and peroxisome to assess localization and the effect (if any) of the probe on colony growth and cell morphology (**Supplemental Figure 1A**). As detailed in the Methods, we selected Sec61-GFP (ER), Vph1-GFP (vacuole), and Pex3-GFP (peroxisome) as bright non-perturbing organelle labels for subsequent experiments.

To assess when and how each organelle is delivered to the buds, we imaged organelle behavior through the cell cycle, illustrated in **Figure 1** for single-budded cells. As discussed further below, the timing and morphology of organelle inheritance was similar in single-budded and multi-budded cells.

**Figure 1:**
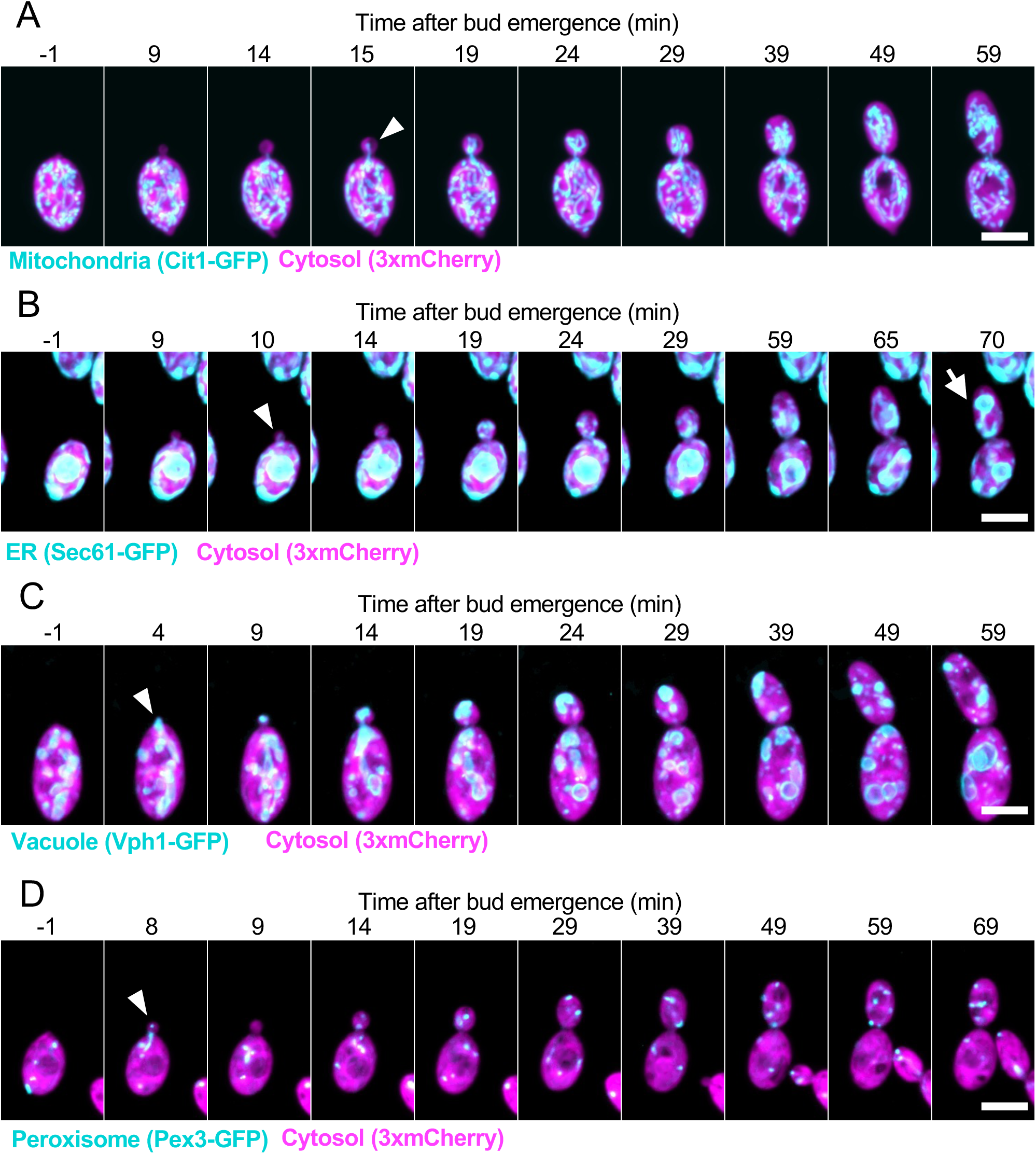
Organelle delivery to buds. Maximum intensity projections of confocal time series of cells expressing a cytosol marker (3xmCherry, magenta) and different organelle markers in cyan: (**A**) mitochondria: Cit1-GFP (DLY25312), (**B**) ER: Sec61-GFP (DLY24944), (**C**) vacuole: Vph1-GFP (DLY24947), and (**D**) peroxisomes: Pex3-GFP (DLY25001). In all panels, white arrowheads indicate when the organelle was first detected in the bud. In **B** the white arrow indicates nuclear envelope entering the bud during mitosis. Note that in **D** a peroxisome is first detected in the bud at minute 8, and in the next minute the peroxisome returned to the mother cell. Scale bar, 5 µm.

Organelle morphology and dynamic behavior was broadly similar to what has been described for *S. cerevisiae* (Rossanese *et al*., 2001; Du *et al*., 2004; Weisman, 2006; Fagarasanu *et al*., 2010; Westermann, 2014; Knoblach and Rachubinski, 2015). Mitochondria formed a tubular network, and they began to populate the bud a few minutes after bud emergence (**Figure 1A**). The ER was a mix of nuclear envelope, tubular elements, and cortical sheets (**Figure 1B, Supplemental Figure 2A**). Cortical ER also entered the bud shortly after bud emergence, while the nuclear envelope remained in the mother until anaphase (Petrucco *et al*., 2024). Vacuoles were ovoid in shape and present in a range of sizes (**Figure 1C**). A portion of a larger mother cell vacuole appeared to be pulled into the bud, followed by fission leaving a smaller portion of the vacuole in the bud. In addition, smaller vacuoles were seen to transit from mother to bud during bud growth. Peroxisomes were present as small puncta whose numbers fluctuated in both mother and bud (**Figure 1D**). Rapid imaging revealed that they traveled bidirectionally between mother and bud, with periods of rapid transit alternating with more stationary behavior (**Supplemental video 1**).

### Inheritance of each organelle follows a stereotyped temporal profile

In what follows, we use the total fluorescence signal of a given organelle probe as a proxy for organelle volume or surface area. We do not distinguish between volume and surface area, as this would require imaging at higher resolution than is possible with light microscopy. However, the organelle geometry for mitochondria (tubes) and ER (thin tubules and sheets) would predict that volume and surface area increase in a linearly correlated manner. Even for vacuoles and peroxisomes, which are more spherical, inheritance involves mainly a change in the number (rather than size) of the organelles present in each bud, so that again the summed volume and surface area would be correlated. In principle, the amount of a fluorescent probe in a bud could reflect the amount of the labeled organelle or a different concentration of probe within the organelle. Visual examination suggested that probe concentration remained uniform, so that the total fluorescence signal of a given organelle probe could be used as a proxy for organelle amount. This was supported by the observation that total fluorescence signal of each probe was well correlated with the total volume of the relevant organelle as measured using 3D segmentation (**Supplemental Figure 2**). In **Figure 2**, we illustrate the patterns of organelle inheritance with example two- or three-budded cells in which the sibling buds behaved similarly.

**Figure 2:**
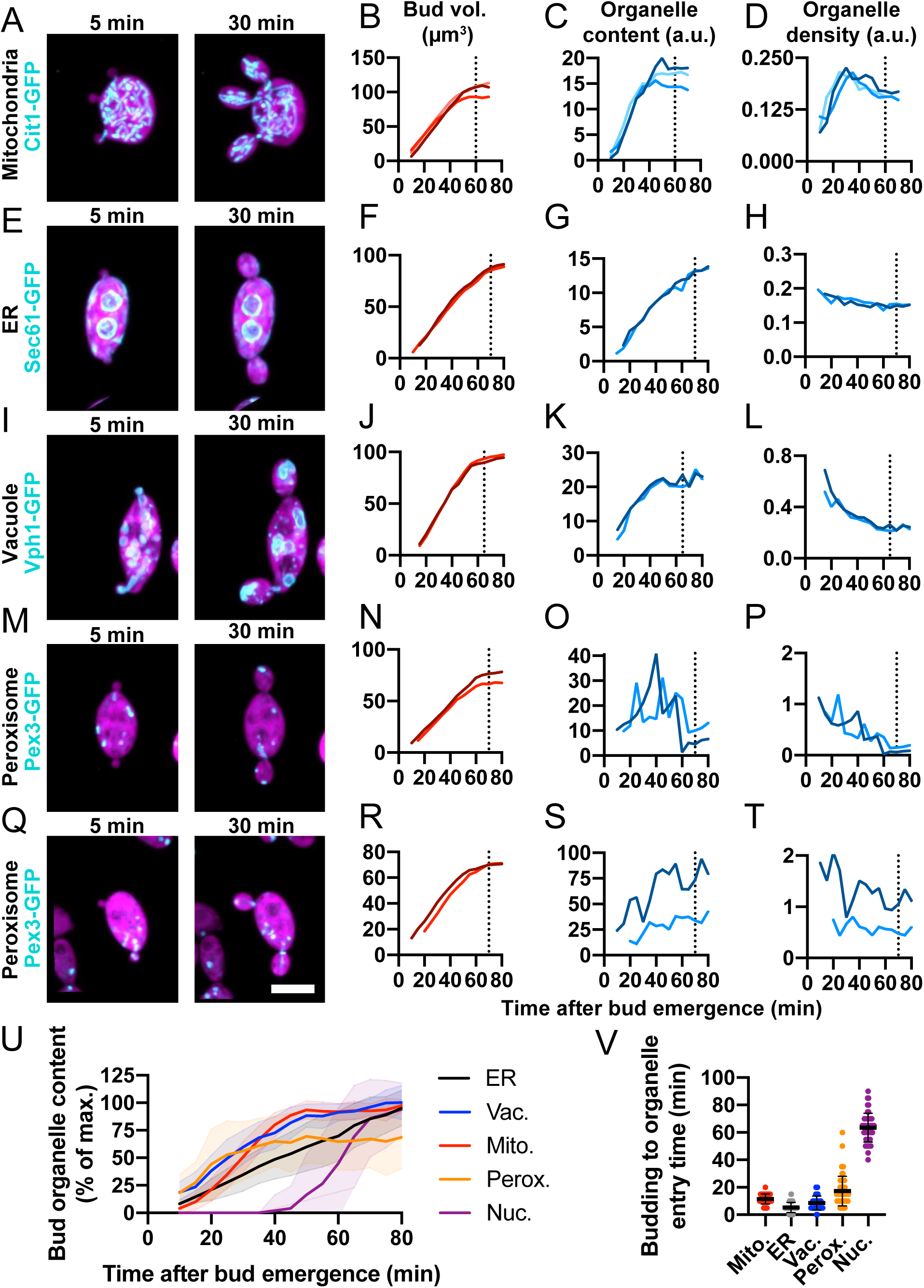
Organelle inheritance follows stereotyped kinetics. Maximum intensity projections of confocal time series of multi-budding cells expressing a cytosol marker (3xmCherry, magenta) and different organelle markers in cyan: (**A**) mitochondria: Cit1-GFP (DLY25312), (**E**) ER: Sec61-GFP (DLY24944), (**I**) vacuole: Vph1-GFP (DLY24947), and (**M** and **Q**) peroxisome: Pex3-GFP (DLY25001). (**B, F, J, N, R**) Quantification of bud growth for the buds shown to the left. (**C, G, K, O, S**) Quantification of total organelle content for the buds shown to the left. (**D, H, L, P, T**) Quantification organelle density (content per bud volume) for the buds shown to the left. In each case the darker and lighter traces refer to different buds from the same mother. (**U**) Mean and standard deviation of total organelle content reported as the percent of the maximum organelle content in each bud over time. n = 42 buds for mitochondria, 49 for ER, 38 for vacuole, and 54 for peroxisome. (**V**) The time between when the bud is first visible (bud emergence) and when the organelle first enters the bud for the same buds quantified in U. The mean and standard deviation are shown. Scale bar, 5 µm.

Mitochondria entered buds 11.4 ± 3.7 min (mean ± s.d., n = 42 buds) after bud emergence (**Figure 2A-C and Supplemental Video 2**). Initially, mitochondrial accumulation outpaced bud growth, such that mitochondrial density (mitochondrial content per bud volume) increased over ∼40-45 min (**Figure 2D**). Then, mitochondrial delivery slowed while buds continued to grow, resulting in a plateau in mitochondrial content (**Figure 2D**).

The cortical ER entered buds 5.3 ± 3.7 min (mean ± s.d., n = 50 buds) after bud emergence (**Figure 2E-G and Supplemental Video 2**). Unlike mitochondria, ER accumulation closely paralleled bud growth, resulting in a constant ER density (**Figure 2H**).

Vacuoles frequently elongated as they entered buds 8.6 ± 5.1 min (mean ± s.d., n = 38 buds) after bud emergence (**Figure 2I-K and Supplemental Video 2**), and then underwent fission, dividing between mother and bud. As buds continued to grow, delivery of small vacuoles continued for ∼40-45 min and then ceased, leading to a dip in vacuole density (**Figure 2L**).

As peroxisome inheritance exhibited greater variability, we provide two illustrative example cells. The timing of peroxisome entry into buds was highly variable, ranging from 5 to 60 min after bud emergence (**Figure 2M-O and Q-S**). Because peroxisomes trafficked bidirectionally between mother and bud, the dynamics of peroxisome content showed large fluctuations (**Figure 2P,T**).

Overall, inheritance of the mitochondria, cortical ER, vacuoles, and peroxisomes began early in the budding cycle and proceeded throughout most of bud growth (**Figure 2U**). This is well before mitosis at 63.5 ± 10.5 min (mean ± s.d., n = 49 buds), when the nucleus enters the bud (**Figure 2V**). Cytokinesis followed at 70 ± 10.4 min (mean ± sd, n = 150 buds). While all organelles were transported into buds, each followed a distinct and stereotyped pattern that was consistent across different buds (**Figure 2U,V**).

### Partitioning of organelles among sibling buds

To ask how evenly the organelles were apportioned among sibling buds in multi-budded cells, we quantified the variability of organelle content (total amount) as well as organelle density (amount per bud volume) among sibling buds. As a measurement of variability, we used the coefficient of variation (CV = standard deviation/mean). A higher CV indicates the parameter is more variable. To avoid noise from very low organelle contents, we only analyzed cells whose buds were larger than 5 µm^3^. This included buds that had recently undergone cytokinesis (detected by a narrow gap in cytoplasmic fluorescence at the neck). To illustrate the cases where inheritance is most variable, we report the relative amounts of organelles in mature buds (volume >50 µm^3^ or just after cytokinesis) from the 10% of cells with the highest CV in organelle density (“outliers”). We also report the frequency with which buds failed to inherit a given organelle altogether.

Mitochondria partitioning to sibling buds exhibited an average CV of 0.28 (n = 133 mother cells), with most siblings varying less than two-fold in total mitochondrial content (**Figure 3A,B**). Much of this variability was clearly attributable to the variability in sibling bud size, as the average CV of mitochondrial density was only 0.16 (n = 133 mother cells) (**Figure 3B**). There was a clear correlation between bud volume and mitochondrial content (**Figure 3C**), even in *pak1Δ* mutants that fail to equalize polarity sites and thus grow buds at different rates producing buds of variable sizes (**Figure 3D,E**) (Crocker *et al*., 2025; Wirshing *et al*., 2025). Fluctuations in mitochondrial density were most marked in small buds, with mature sibling buds having a smaller CV (**Figure 3F**). In the outliers, siblings differed on average 2.6-fold in mitochondrial density (n = 78 mother cells with mature buds) (**Figure 3G**). Out of 1323 growing buds monitored by time lapse microscopy, all received mitochondria. Thus, mitochondrial delivery is generally well correlated with bud growth, differing only slightly between sibling buds, and is robust with all buds scored receiving mitochondria.

**Figure 3:**
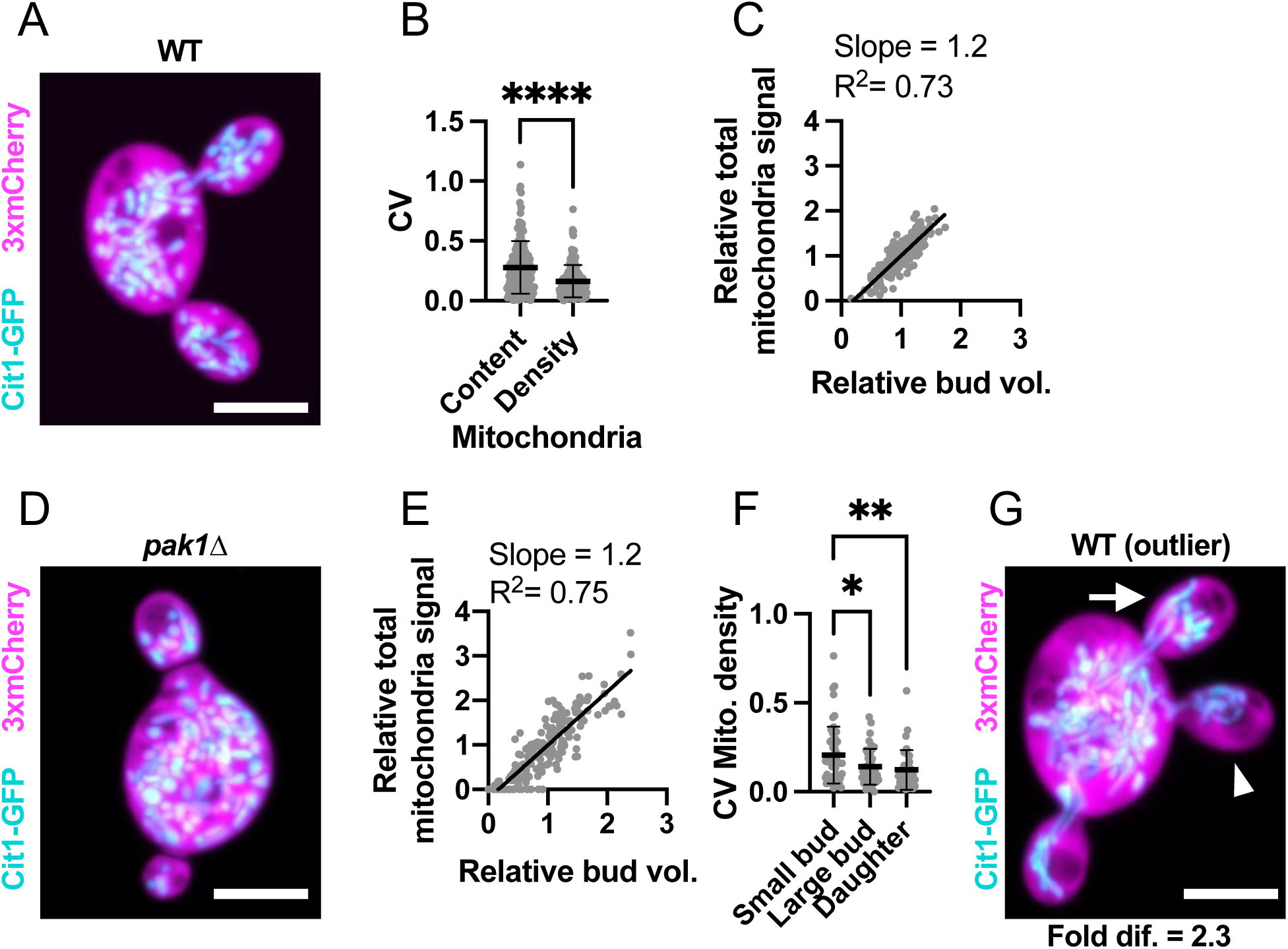
Mitochondria are evenly partitioned among sibling buds. (**A**) Maximum intensity projection of a confocal Z-series of a wild-type cell expressing a mitochondrial marker (cyan, Cit1-GFP) and cytosol marker (3xmCherry, magenta) (DLY25312). (**B**) Variability (CV) of sibling mitochondrial content (Mito.) and mitochondria content per bud volume (Mito./Bud vol.). The mean and standard deviation are shown. ****, p ≤ 0.0001 by two-tailed Student’s t-test; n = 133 mother cells. (**C**) Relationship between bud volume and total mitochondrial content, both relative to siblings, in wild-type cells. The best-fit slope and R^2^ (square of Pearson correlation coefficient) are shown (n = 140 buds). (**D**) Image (as in A) of a *pak1Δ* cell with differently sized buds (DLY26344). (**E**) Relationship between bud volume and mitochondrial content (as in C) in *pak1Δ* cells. n = 180 buds. (**F**) Variability (CV) of sibling mitochondria content per bud volume in small buds (sm. buds, <50 µm^3^), large buds (lg. buds, >50 µm^3^), and daughter cells just after cytokinesis (daughters) (n = 55, 41, and 37, respectively). The mean and standard deviation are shown. Statistical significance calculated by one-way ANOVA (not shown, p > 0.05; *, p ≤ 0.05; **, p ≤ 0.01). (**G**) Image (as in A) of an example outlier wild-type cell (DLY25312). The arrowhead indicates the bud with the least mitochondria and the arrow the bud with the most. The fold difference between these buds is indicated. Scale bar, 5 µm.

Cortical ER partitioning to sibling buds exhibited a similar average CV to that of mitochondria (0.20, n = 156) (**Figure 4A,B**), and again there was a clear correlation between bud volume and cortical ER content in both wild-type and *pak1Δ* buds (**Figure 4C-E**). Unlike mitochondria, ER density was consistent among siblings in both smaller and mature buds (**Figure 4F**). Even in the outliers, siblings differed only 1.8-fold on average in cortical ER density (n = 125 mother cells with mature buds) (**Figure 4G**). Some of this variability appeared to be due to buds inheriting different numbers of nuclei and therefore different amounts of ER associated with the nuclear envelope (**Figure 4G**). However, siblings could also differ in cortical ER content prior to mitosis (**Figure 4G**). Of the 1441 growing buds monitored, all received ER. Consistent with our observation that cortical ER enters buds much earlier than nuclei (**Figure 2U,V**), even a very rare bud that failed to inherit nuclei still inherited cortical ER. Thus, buds accurately inherit ER in proportion to their size.

**Figure 4:**
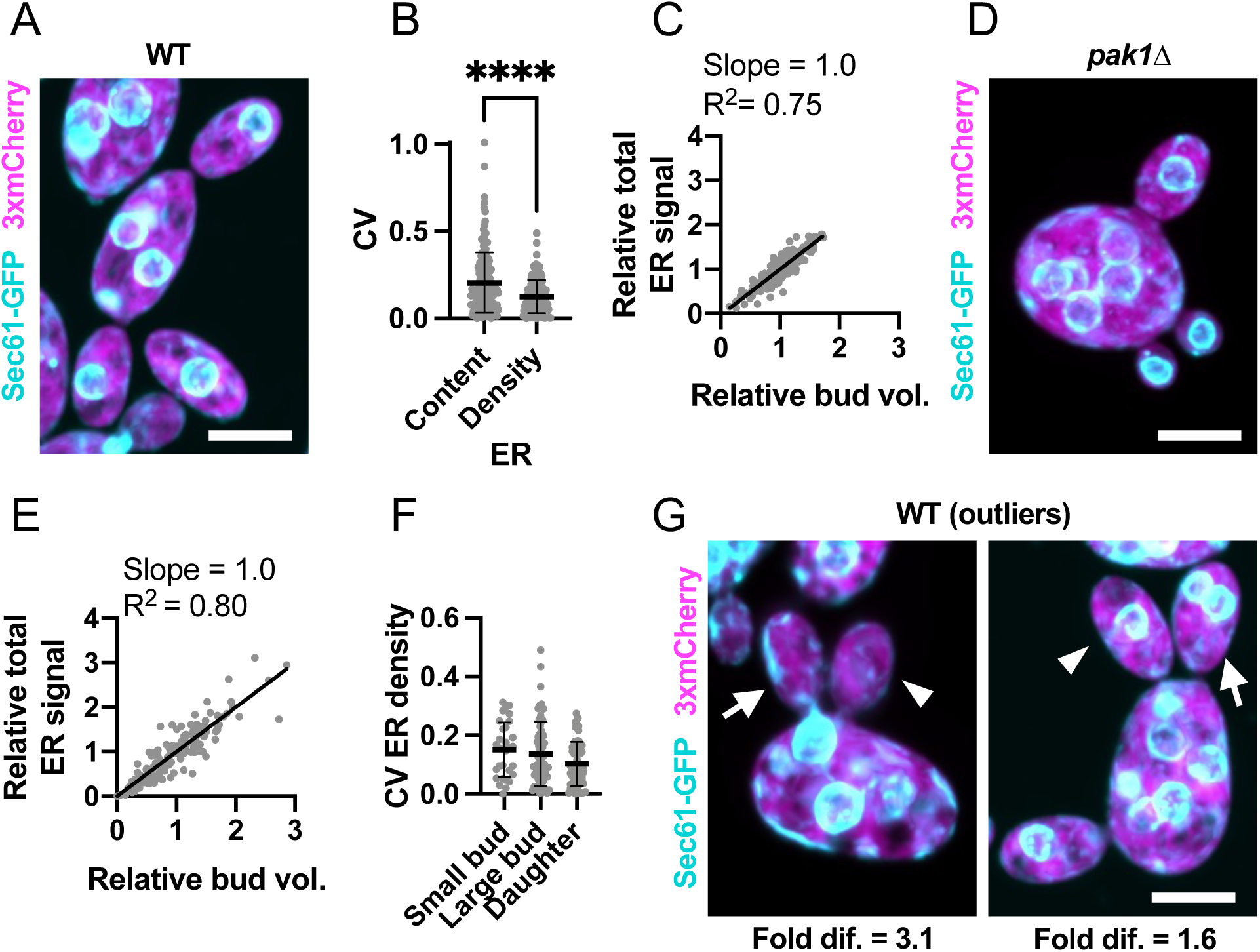
The ER is evenly partitioned among sibling buds. (**A**) Maximum intensity projection of a confocal Z-series of a wild-type cell expressing an ER marker (cyan, Sec61-GFP) and cytosol marker (3xmCherry, magenta) (DLY24944). (**B**) Variability (CV) of sibling ER content (ER) and ER content per bud volume (ER/Bud vol.). The mean and standard deviation are shown. ****, p ≤ 0.0001 by two-tailed Student’s t-test; n = 156 mother cells. (**C**) Relationship between bud volume and total ER content, both relative to siblings, in wild-type cells. The best-fit slope and R^2^ (square of Pearson correlation coefficient) are shown (n = 396 buds). (**D**) Image (as in A) of a *pak1Δ* cell with differently sized buds (DLY26341). (**E**) Relationship between bud volume and ER content (as in C) in *pak1Δ* cells. n = 153 buds. (**F**) Variability (CV) of sibling ER content per bud volume in small buds (sm. buds, <50 µm^3^), large buds (lg. buds, >50 µm^3^), and daughter cells just after cytokinesis (daughters) (n = 31, 60, and 65, respectively). The mean and standard deviation are shown. p > 0.05 by one-way ANOVA. (**G**) Images (as in A) of example outlier wild-type cells (DLY24944). The arrowhead indicates the bud with the least ER and the arrow the bud with the most. The fold difference between these buds is indicated. Scale bar, 5 µm.

Vacuole partitioning to sibling buds exhibited an average CV of 0.42 (n = 174 mother cells) (**Figure 5A,B**), significantly higher than the CVs for mitochondrial and ER inheritance. Moreover, bud volume and vacuole content were only poorly correlated (**Figure 5C-E**). This variability decreased only slightly in mature sibling buds (**Figure 5F**). Outliers had sibling buds that differed on average 8.4-fold in vacuole density (n = 104 mother cells with mature buds) (**Figure 5G**). Despite this variability, out of 1421 growing buds monitored, all received vacuoles. Thus, unlike ER and mitochondria, vacuole inheritance is not proportional to bud size and can differ between siblings. However, as with the ER and mitochondria, mother cells ensure each daughter inherits at least one vacuole.

**Figure 5:**
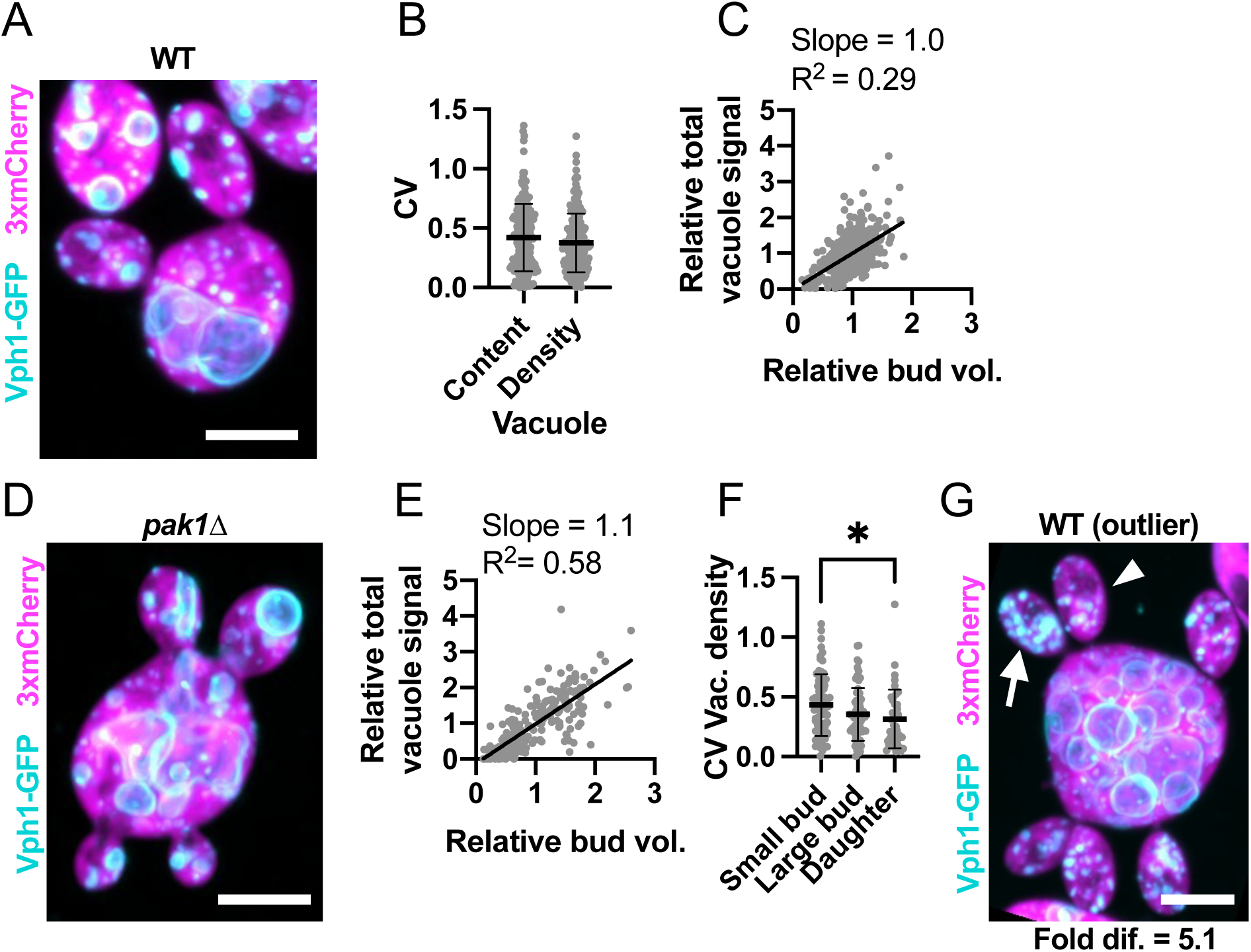
Vacuole inheritance among sibling buds is variable. (**A**) Maximum intensity projection of a confocal Z-series of a wild-type cell expressing a vacuole marker (cyan, Vph1-GFP) and cytosol marker (3xmCherry, magenta) (DLY24947). (**B**) Variability (CV) of sibling vacuole content (Vac.) and vacuole content per bud volume (Vac./Bud vol.). The mean and standard deviation are shown. p > 0.05 by two-tailed Student’s t-test. n = 174 mother cells. (**C**) Relationship between bud volume and vacuole content, both relative to siblings, in wild-type cells. The best-fit slope and R^2^ (square of Pearson correlation coefficient) are shown (n = 532 buds). (**D**) Image (as in A) of a *pak1Δ* cell with differently sized buds (DLY26335). (**E**) Relationship between bud volume and vacuole content (as in C) in *pak1Δ* cells (n = 199 buds). (**F**) Variability (CV) of sibling vacuole content per bud volume in small buds (sm. buds, <50 µm^3^), large buds (lg. buds, >50 µm^3^), and daughter cells just after cytokinesis (daughters) (n = 70, 63, and 41, respectively). The mean and standard deviation are shown. Statistical significance calculated by one-way ANOVA (not shown, p > 0.05; *, p ≤ 0.05). (**G**) Image (as in A) of an example outlier wild-type cell (DLY24947). The arrowhead indicates the bud with the least vacuole and the arrow the bud with the most. The fold difference between these buds is indicated. Scale bar, 5 µm.

Peroxisome partitioning was even more variable, with an average CV of 0.69 (n = 173 mother cells) (**Figure 6A,B**). This variability was not due to variation in bud volume, as peroxisome content was not correlated with bud volume in WT or *pak1Δ* cells (**Figure 6C-E**). Newborn daughters that had recently completed cytokinesis were only slightly less variable than small buds (**Figure 6F**). In the outliers, siblings differed by an astonishing 23.2-fold (on average) in peroxisome content per volume (**Figure 6G**). Additionally, daughter cells sometimes failed to inherit peroxisomes. Of the 1289 buds monitored, 20 did not contain peroxisomes at the time of cytokinesis (**Supplemental Video 3**). We also observed instances where mother cells failed to retain peroxisomes (**Supplemental Video 3**). In mothers or buds without peroxisomes, new Pex3-GFP puncta appeared 30-60 min after cytokinesis, suggesting that *de novo* peroxisome synthesis took place (**Supplemental Video 3**). Thus, unlike the other organelles, mother cells do not ensure that each bud inherits a peroxisome, and peroxisome inheritance is highly variable.

**Figure 6:**
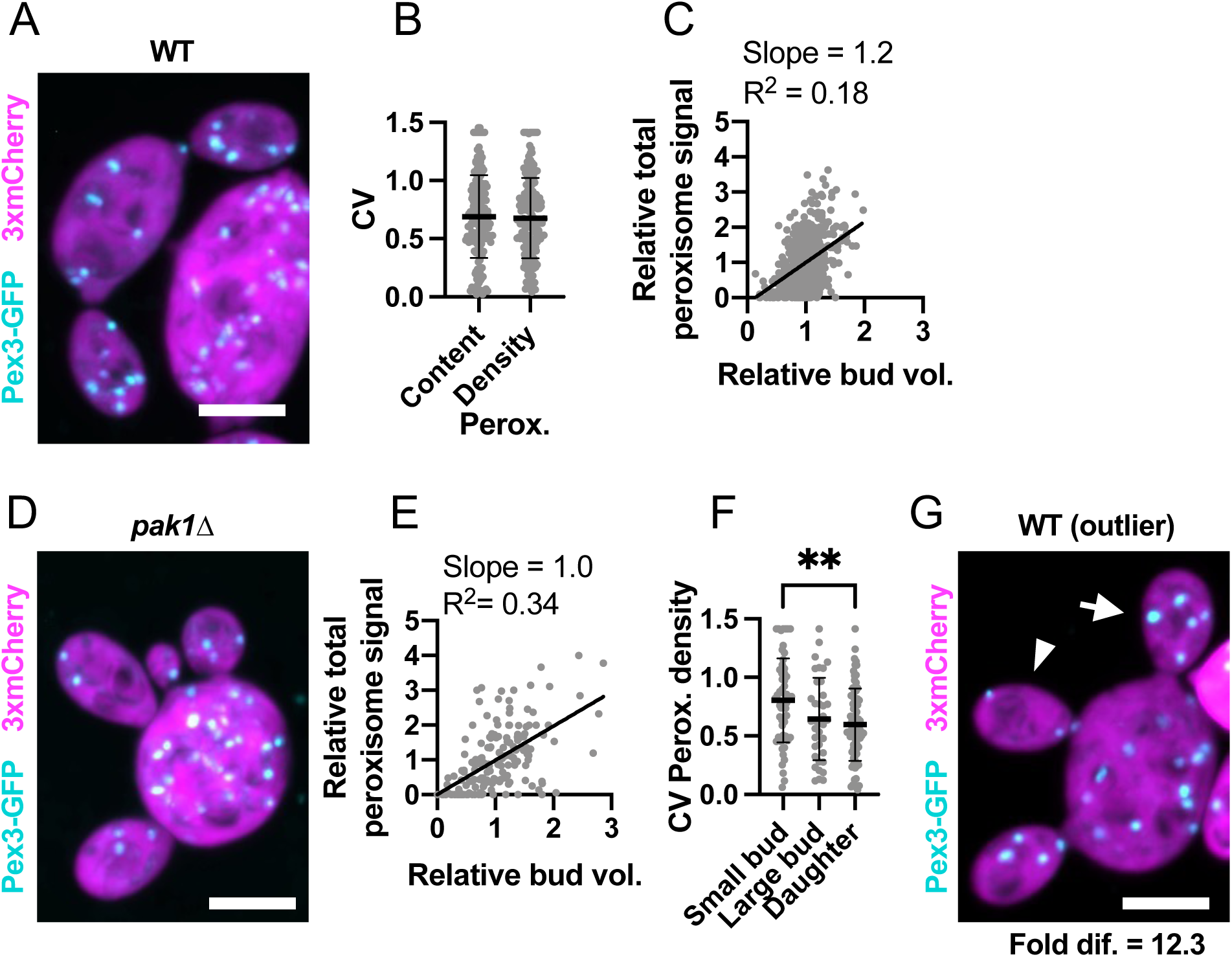
Peroxisome partitioning among sibling buds is highly variable. (**A**) Maximum intensity projection of a confocal Z-series of a wild-type cell expressing a peroxisome marker (cyan, Pex3-GFP) and cytosol marker (3xmCherry, magenta) (DLY25001). (**B**) Variability of sibling peroxisome content (Perox.) and peroxisome content per bud volume (Perox./Bud vol.). The CV = coefficient of variation. The mean and standard deviation are shown. p > 0.05 by two-tailed Student’s t-test. n = 173 mother cells. (**C**) Relationship between bud volume and peroxisome content, both relative to siblings, in wild-type cells. The best-fit slope and R^2^ (square of Pearson correlation coefficient) are shown (n = 657 buds). (**D**) Image (as in A) of a *pak1Δ* cell with differently sized buds (DLY26338). (**E**) Relationship between bud volume and peroxisome content (as in C) in *pak1Δ* cells (n = 175 buds). (**F**) Variability (CV) of sibling peroxisome content per bud volume in small buds (sm. buds, <50 µm^3^), large buds (lg. buds, >50 µm^3^), and daughter cells just after cytokinesis (daughters) (n = 58, 37, and 78, respectively). The mean and standard deviation are shown. Statistical significance calculated by one-way ANOVA (not shown, p > 0.05; **, p ≤ 0.01). (**G**) Image (as in A) of an example outlier wild-type cell (DLY25001). The arrowhead indicates the bud with the fewest peroxisomes and the arrow the bud with the most. The fold difference between these buds is indicated. Scale bar, 5 µm.

### Uneven organelle partitioning is largely stochastic and organelle-specific

The analysis above shows that outliers, defined as the 10% of mothers with highest variability in organelle partitioning among their buds, can bequeath uneven complements of any given organelle to their buds. However, looking at only one organelle at a time, it is not possible to assess whether uneven inheritance is correlated between organelles. At one extreme, uneven inheritance might be the product of stochastic errors that only affect a single organelle, in which case an outlier in vacuole inheritance might deliver equal amounts of mitochondria to each bud, and so on. Alternatively, there might be a subset of “hyper-variable mothers” that unequally partition all organelles. It is also possible that unequal inheritance of one organelle might causally impact the inheritance of other organelles. For example, a bud that received an unusually large vacuole may have insufficient room to receive the normal complements of other organelles.

To ask whether uneven organelle partitioning exhibits stochastic or correlated features, we imaged a strain with three compatible fluorescent organelle probes: Vph1-3xGFP for vacuoles, Sec61-3xmCherry for ER, and a far-red MitoTracker dye for mitochondria (see Methods) (**Figure 7A and Supplemental Video 4**). In this experiment, we were able to assess whether heterogeneity in organelle partitioning was correlated among organelles. We found no correlation between the CVs of mitochondrial, ER, and vacuole inheritance (**Figure 7B-D**). Thus, uneven inheritance is not a consequence of systematic differences among mother cells in their ability to partition delivery. We also assessed whether the inherited amounts of the different organelles were correlated. However, buds receiving more or less of one organelle did not systematically receive more or less of a different organelle (**Figure 7E-F**). Rather, each organelle exhibited uncorrelated differences in partitioning, suggesting that uneven inheritance is a result of organelle-specific stochastic differences.

**Figure 7:**
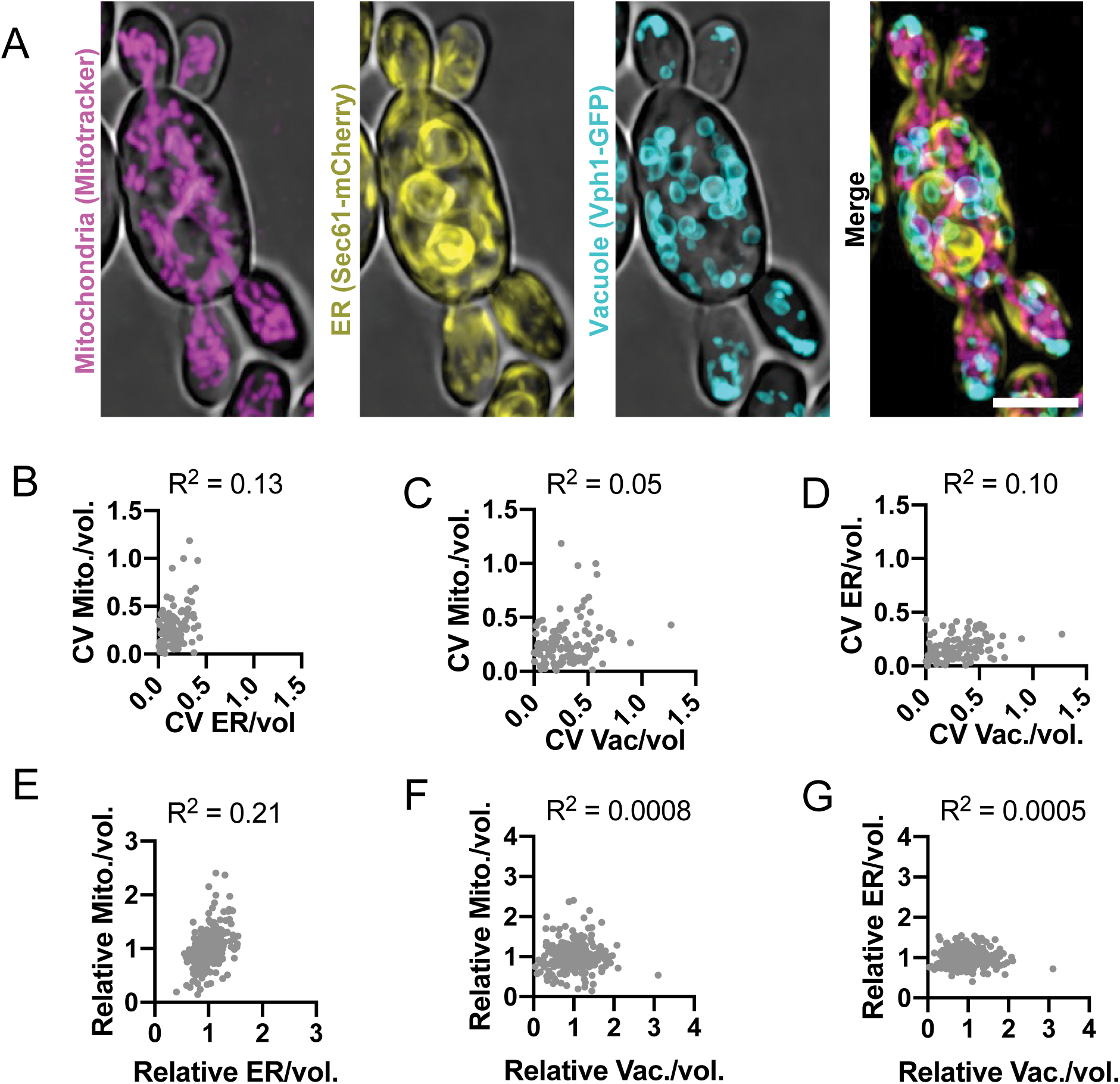
Inheritance variability is uncorrelated across different organelles. (**A**) Maximum intensity projection of a confocal Z-series of a wild-type cell stained with a mitochondria stain (magenta, MitoTracker) and expressing an ER marker (yellow, Sec61-3xmCherry) and vacuole marker (cyan, Vph1-3xmNG) (DLY26775). Scale bar, 5 µm. (**B**) Relationship between ER density variability (CV) and mitochondrial density variability, both relative to sibling buds (n = 111 mother cells). R^2^, square of Pearson correlation coefficient. (**C**) Relationship between vacuole content variability and mitochondrial content variability for the same cells analyzed in B. (**D**) Relationship between vacuole content variability and ER content variability for the same cells analyzed in B and D. (**E**) Relationship between a bud’s ER content and mitochondria content (n = 366 buds). R^2^, square of Pearson correlation coefficient. (**F**) Relationship between a bud’s vacuole content and mitochondrial content for the same buds analyzed in E. (**G**) Relationship between a bud’s vacuole content and ER content for the same buds analyzed in E.

### Organelle variability among newly born daughter cells from different mothers

In our analysis thus far, we focused on the variability in organelle inheritance between siblings from the same mother. However, it seemed possible that there might be a second layer of variability stemming from differences between mother cells in terms of how much of a given organelle they retain versus how much they deliver to their buds. In a similar vein, previous work indicated that the size variation among newborn daughters in the population was significantly greater than the size variation among siblings, because mother cells make variable numbers of buds (Wirshing *et al*., 2025). If mother cells similarly differ in their propensity to deliver organelles, then the CV in organelle density (content per cell volume) across an entire population of newborn daughters might exceed the CV among sibling daughters.

We used the same strains carrying individual organelle markers to quantify organelle density in mature buds (> 50 µm^3^) and daughter cells just after cytokinesis. This analysis showed that the CV in organelle content across the entire population was somewhat higher than the CV for the same organelle among siblings (**Figure 8A**). Thus, variability between mothers as well as between siblings contributes to population variability. Comparing normalized organelle densities (normalized to the population mean for mature buds/newborn cells) revealed the same trends in variability as we documented for sibling buds (**Figure 8B**). Thus, population variability stems in large part from uneven inheritance. To quantify this variability in organelle content in another way, we also report the fold-difference in organelle density between the 10% of cells with highest and lowest organelle densities (**Figure 8C**). This analysis lays the foundation for future studies to examine the consequences of organelle variability.

**Figure 8:**
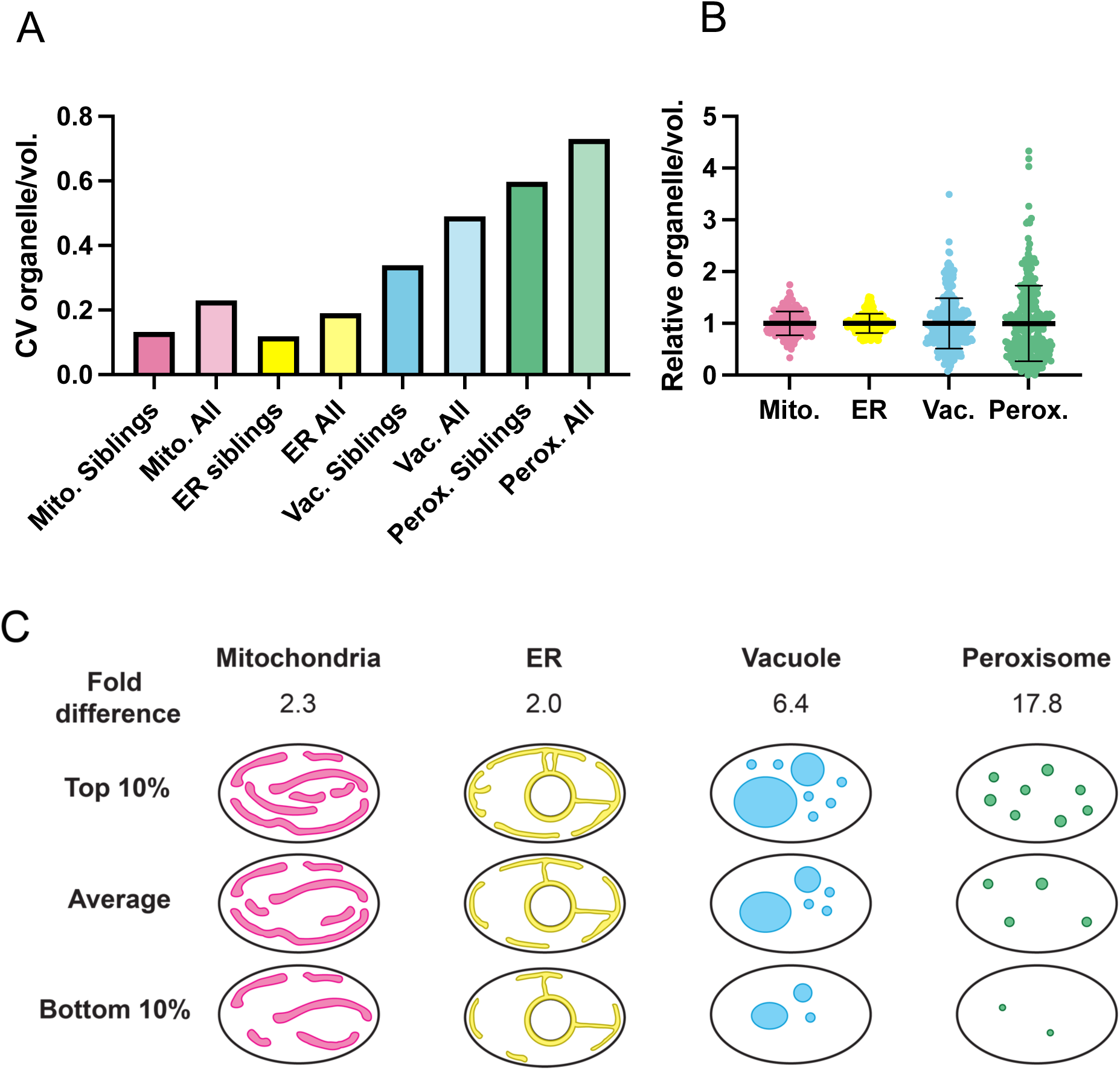
Organelle variability in the population of daughter cells. (**A**) CV in organelle density in mature buds (>50 µm^3^) and newborn daughters, comparing data from sibling buds (derived from Fig.s 3-6) and from the entire population (n = 198, 219, 326, and 272 buds or daughters for mitochondria, ER, vacuole, and peroxisome, respectively). (**B**) Organelle density variation across the entire population of mature buds and newborn daughters of the same cells used to measure the population CV in B. The mean and standard deviation are shown. (**C**) A schematic illustrating organelle content variability across the population with the fold difference in organelle density in the top vs bottom 10% of cells indicated.

## DISCUSSION

Stochastic molecular-level differences in protein copy number (noise) can produce significant cell-to-cell variation in phenotype, particularly for small cells (Maheshri and O’Shea, 2007; Losick and Desplan, 2008; Raj and Van Oudenaarden, 2008; Eldar and Elowitz, 2010). Such stochasticity can be deleterious, as when cells lack sufficient amounts of a key molecule, or beneficial, as when stochasticity enables bet-hedging strategies. As low copy-number cell constituents, organelle numbers also have the potential to drive stochastic differences between genetically identical cells. Thus, variation in organelle content has the potential to produce considerable phenotypic variation even in large cells. It is clear that failure to inherit a key organelle like the nucleus or mitochondria would be catastrophic for the cell, but the effects of less extreme variation in copy number are not as clear. Unlike the case of gene expression noise, we are only beginning to appreciate the natural degree of variation in organelle composition in some model systems like *S. cerevisiae* (Uchida *et al*., 2011; Rafelski *et al*., 2012; Mukherji and O’Shea, 2014; Chan *et al*., 2016). Here, we provide a quantitative analysis of organelle inheritance in a multi-budding yeast. Our findings suggest that variability in organelle inheritance is tuned to different levels for different organelles, suggesting the existence of mechanisms to reduce stochasticity in ER and mitochondrial inheritance, while allowing much more variation in vacuole and peroxisome inheritance. Consistent with these findings, mitochondrial content appeared to be less variable than vacuole content in *S. cerevisiae* (Uchida *et al*., 2011). Interestingly, vacuole and peroxisomes can be made *de novo* (Weisman *et al*., 1990; Hoepfner *et al*., 2005), whereas ER and mitochondria cannot.

### Inheritance of cortical ER and mitochondria

In *S. cerevisiae*, bud growth and organelle delivery are both directed by a polarized network of actin cables (Knoblach and Rachubinski, 2015). Actin cables are nucleated at the growing bud tip and serve as tracks for myosin V transport of multiple organelles (including all of those analyzed here) as well as post-Golgi secretory vesicles that drive bud growth (Knoblach and Rachubinski, 2015). Actin cables similarly direct bud growth in *A. pullulans*, but in this system multiple polarity sites can be established simultaneously (Crocker *et al*., 2025). Prior to bud emergence, polarity sites are equalized, leading to the assembly of similar actin cable networks that deliver vesicles to each bud at similar rates (Wirshing *et al*., 2025). Disrupting the cell’s ability to equalize polarity sites (as in *pak1Δ* mutants) yields sibling buds that grow at different rates and reach different sizes (Crocker *et al*., 2025; Wirshing *et al*., 2025). Here we find that in both wild-type and *pak1Δ* cells, cortical ER and mitochondria are inherited in a manner that tracks closely with bud growth. Thus, the simplest way to explain our findings is that as in *S. cerevisiae*, the same actin cable networks that direct the delivery of vesicles also deliver ER and mitochondria.

### Vacuole and peroxisome inheritance

The morphology and timing of vacuole delivery in *A. pullulans* was very similar to that described for *S. cerevisiae*, suggesting that vacuoles too are delivered by the same actin cable networks. However, partitioning of the vacuoles among sibling buds was more variable, and showed much less correlation between bud size and vacuole content. Thus, either vacuole delivery on the same tracks is more variable, or vacuole inheritance is affected by other factors. Like vacuoles, peroxisome inheritance was highly variable between sibling buds, and the amount inherited was uncorrelated with bud size. In this case, however, the dynamics of peroxisome inheritance were unlike those in *S. cerevisiae*, with bidirectional movements of peroxisomes between mother and buds.

While *S. cerevisiae* cells lack an interphase network of microtubules (Adams and Pringle, 1984; Kilmartin and Adams, 1984), *A. pullulans* has an extensive interphase microtubule network (Petrucco *et al*., 2025). Microtubules are not required to drive bud growth (Wirshing *et al*., 2025), but could play a role in organelle inheritance. Many other fungi use interphase microtubule networks to position organelles (Herr and Heath, 1982; Yaffe *et al*., 1996; Steinberg *et al*., 1998; Wu *et al*., 1998; Seiler *et al*., 1999; Steinberg, 2015). Our observations on peroxisomes are reminiscent of studies in *A. nidulans* and *U. maydis*, where peroxisomes move rapidly and bidirectionally by hitchhiking on early endosomes traveling along microtubules (Guimaraes *et al*., 2015; Salogiannis *et al*., 2016). Thus, it is possible that the increased variability we observed for vacuoles and peroxisomes in *A. pullulans* is due to microtubule-based transport.

For all organelles except peroxisomes, inheritance mechanisms appear to be very robust, in that none of the daughters we observed were born without ER, vacuoles, or mitochondria (n > 1,300 in each case). Mitotic nuclear delivery is similarly robust, although occasional errors can lead to anucleate buds (∼1 in 2,000 buds) (Petrucco et al. 2025). In contrast, one in every ∼64 newborn daughters (and also some mothers) did not inherit peroxisomes. In these cases, de novo peroxisome synthesis later restored the organelle.

### Open questions

Our findings raise several questions. Given that daughters can inherit quite different organelle contents, what are the downstream phenotypic effects of such differences? Are the differences compounded in subsequent generations? Or do organelle biogenesis pathways return the organelle distributions to a more uniform state?

If actin- and microtubule-based transport systems collaborate in organelle partitioning, how is traffic by these distinct networks is coordinated during budding growth? Is microtubule-mediated inheritance intrinsically more variable? If so, is such variability desirable for some organelles? Why would that be the case?

Why are vacuole and peroxisome inheritance so much more variable than ER or mitochondria inheritance? The need for processes like autophagy (requiring vacuoles) or fatty acid β-oxidation (occurring in peroxisomes) varies greatly depending on the environment (Veenhuis *et al*., 1987; Valenciano *et al*., 1996; Lei *et al*., 2022). If stressors that create a requirement for more vacuoles or peroxisomes can arise rapidly in changing environments, then variable inheritance of vacuole or peroxisomes may constitute a bet-hedging strategy to ensure that some progeny are well prepared for any eventuality.

Our focus in this report has been on the inheritance of organelles by newborn daughters, which tend to be the smallest cells in a population of *A. pullulans*, and perhaps serve as vehicles for dispersal in marine environments (Kurita *et al*., 2026). However, budding is not the only option for these cells, which can also undergo cell cycles in which mother cells enlarge without budding and without cytokinesis (Petrucco *et al*., 2025). As a result, mother cells can vary greatly in volume and number of nuclei. In the accompanying report we ask how organelles scale with cell size in a proliferating population whose cells can vary over a remarkable 100-fold size range.

## METHODS

### *A. pullulans* strains and maintenance

All strains used in this study are in the *Aureobasidium pullulans* EXF-150 (Gostinčar *et al*., 2014) strain background, with genetic modifications listed in **Supplemental Table 1**. Unless otherwise indicated, *A. pullulans* was grown at 24°C in standard YPD medium (2% glucose, 2% peptone, 1% yeast extract) with 2% BD Bacto^TM^ agar (214050, VWR) in plates.

### Strain construction

Organelle markers were introduced into a strain expressing the cytoplasmic marker 3xmCherry (DLY24919) described previously (Wirshing *et al*., 2025). C-terminal GFP or mNG tags were introduced at the native loci of genes encoding vacuole-localized proteins Atg42 (protein ID 347152) and Vph1 (protein ID 348720), ER-localized proteins Sec61 (protein ID 345851), Elo3 (protein ID 363594), and May24 (protein ID 375917), and peroxisome-localized proteins Pex3 (protein ID 330605), Pex14 (protein ID 402714), Pex5 (protein ID 282059), and Pex11 (protein ID 395491). Mitochondria were visualized by C-terminally tagging Cit1 with GFP as described previously (Wirshing *et al*., 2024). Integration was targeted to native loci by 3-part-PCR and pAPInt vectors with GFP or three tandem copies of mNG (3xmNG) and the hygromycin or nourseothricin resistance cassettes as described previously (Colarusso *et al*., 2025). Briefly, ∼1000 bp upstream and downstream of the desired integration site and the integration cassette (fluorescent tag + resistance cassette) were all amplified using Phusion™ Hot Start Flex 2X Master Mix (M0536L, NEB) following the manufacturer’s instructions. Primers were designed to introduce 60-70 bp overlaps between the three PCR products. Eight PCRs (50 µl each) were pooled, purified, and concentrated by adding 0.1 volume of 3 M NaOAc and 2.5 volumes of 100% ethanol followed by incubation on ice for 10 min. The DNA-ethanol mixture was then added to a silica DNA binding column (T1017-2, New England Biolabs), rinsed with 400 µl wash buffer (80% EtOH, 20 mM NaCl, 2 mM Tris pH 8), allowed to dry for 1 min, and then eluted in ∼40-60 µl 10 mM Tris pH 8 at a final concentration of ∼1,500 ng/µl.

*A. pullulans* was transformed using PEG/LiOAc/ssDNA as described previously (Wirshing *et al*., 2024). The three separate PCR products were added directly to competent cells during transformation in equimolar ratio (∼15 µl total volume of purified PCR products). After transformation and recovery (4-18 hours at 24°C in YPD + 1M sorbitol with agitation at 100 rpm), cells were selected on YPD plates supplemented with Hygromycin B (400051-1MJ, Millipore) at a final concentration of 174 µl/l (∼70.4 mg/l) or 50 mg/l Nourseothricin (N1200-1.0, Research Products International). Colonies appeared after 2-3 days 24°C. Integration was confirmed by colony PCR using the Phire Plant Direct PCR Master Mix (#F-160S, Thermo Scientific) following the manufacturer’s instructions. All colonies that were positive by colony PCR were also positive for GFP or mNG expression when checked via fluorescence microscopy (see below for imaging conditions). At least three transformants were derived for each genotype, and were confirmed to display qualitatively similar growth rates (**Supplemental Figure 1B**) and fluorescence patterns.

### A. pullulans growth assay

To compare cell growth of *A. pullulans* strains expressing different organelle markers, a single colony was inoculated into 5 ml of YPD (2% glucose) and grown at 24°C for 48 h. Cultures were serially diluted 10-fold five times in sterile deionized water in a 96-well plate and transferred onto solid YPD plates using a pin-frogger. Plates were grown for 48 h at 24°C or 30°C and imaged on an Amersham Imager 680 (General Electric Company) using the colorimetric Epi-white settings. None of the strains expressing organelle markers had detectable growth defects compared to WT at 24°C or 30°C.

### Live-cell imaging and image analysis

To grow yeast for imaging experiments, a single colony was used to inoculate 5 ml of YPD (2% glucose). Cultures were grown overnight at 24°C to a density of 1-5×10^6^ cells/ml. Cells were pelleted at 9391 rcf for 10 s and the supernatant was removed leaving behind enough liquid to resuspended the cell pellet at a final density of ∼7×10^7^ cells/ml. 2 µl of the resuspended cells were added to the bottom of a glass-bottomed 8-well Ibidi chamber (80827, Ibidi) and covered with a 200 mg 5% agarose (97062-250, VWR) pad made with CSM (6.71 g/L BD Difco^TM^ Yeast Nitrogen Base without Amino Acids, BD291940, FisherScientific, 0.79 g/L Complete Supplement Mixture, 1001-010, Sunrise Science Products, and 2% glucose). All experiments were conducted at room temperature (20-22°C).

Cells were imaged on a Nikon Ti2E inverted microscope with a CSU-W1 spinning-disk head (Yokogawa), CFI60 Plan Apochromat Lambda D 60x Oil Immersion Objective (NA 1.42; Nikon Instruments), a LUN-F compact four-line laser source (405 nm 100 mW, 488 nm 100 mW, 561 nm 100 mW, and 640 nm 75 mW fiber output), and a Hamamatsu ORCA Quest qCMOS camera controlled by NIS-Elements software (Nikon Instruments). For still images, the entire cell volume was acquired using 79 Z-slices at 0.2 μm step intervals. Exposure times of 50 ms at 50% laser power were used with excitation at 488 nm (with ET525/36m Emission Filter) to excite GFP or mNG, with excitation at 561 nm (with ET605/52m Emission Filter) to excite mCherry, and with excitation at 640 nm (with ET705/72m Emission Filter) to excite MitoTracker Deep Red FM (M22426, Thermofisher). To track bud growth and organelle inheritance over time, Z-stacks (17 slices, 0.7 µm interval) were acquired every 1-5 minutes for 2-3 hours using 50 ms exposures at 15% laser power for excitation 488 nm and 10% laser power for excitation at 561 nm. To capture rapid peroxisome dynamics, a single Z-slice was acquired every 200 ms for 20 s using 15% laser power with excitation 488 nm to image Pex3-GFP. To increase acquisition speed, mCherry signal in the cytoplasm (which changes little in 20 s) was only imaged in the first frame of these time series using 10% laser power and excitation at 561 nm.

To characterize the probes used to detect each organelle, confocal Z-stacks of each organelle marker strain were denoised and maximum intensity projected using NIS-Elements General Analysis 3 (GA3, Nikon Instruments) software. Vacuole markers Vph1-GFP and Atg42-3xmNG appeared similar and Vph1 was brighter, so this tag was selected for further analysis. ER markers Sec61-GFP, Elo3-GFP, and May24-3xmNG appeared similar but May24-3xmNG was quite dim. Elo3-GFP was enriched in the nuclear envelope, making the peripheral ER more difficult to detect, so we selected Sec61-GFP for further analysis. Peroxisome markers Pex11-GFP and Pex5-3xmNG both resulted in small numbers of large puncta, potentially indicating aggregation or disruption of peroxisome fission, so these markers were excluded from further analysis. Peroxisomes in strains expressing Pex3-GFP or Pex14-2xmNG appeared similar, and we selected Pex3-GFP for further analysis.

To track bud growth and organelle inheritance captured in time lapse experiments, the cytoplasmic 3xmCherry signal was used to 3D segment growing buds for bud volume measurements and the GFP-tagged organelle signal was used to 3D segment each organelle in NIS-Elements General Analysis 3 (GA3, Nikon Instruments) software. We recognize that segmentation of objects below the resolution limit would yield inaccurate volumes, but the process is useful to estimate the correlation between fluorescent intensity and organelle amount (**Supplemental Figure 2**) and to allow subtraction of the cytoplasmic background. After segmentation, GFP signal outside of the organelle mask in the cytoplasm was removed setting the GFP intensity in the cytoplasm to zero. After subtraction of the cytoplasmic background signal, the bud volume and total GFP signal (total organelle content) over time was measured and is reported. To determine when organelles first enter growing buds, the first frame where the GFP signal in the bud was above zero (after cytoplasmic background subtraction) was recorded. A similar approach was used to measure total organelle content (total GFP signal), total organelle volume, and bud volumes in still images. These measurements were used to test for correlations between bud volumes and total organelle content, quantify inheritance variability across siblings and across the whole population of buds and daughter cells.

### Staining mitochondria with MitoTracker

Cells were grown in YPD as described above for imaging experiments. 1.5 ml overnight culture was transferred to a fresh glass culture tube and 0.6 µl of 1 mM MitoTracker Deep Red FM (M22426, Thermofisher) in DMSO was added (final concentration 400 nM). Cells were incubated with the dye for 15 min at 24°C with agitation. Cells were then rinsed two times with YPD and mounted for imaging as described above.

### Statistical analysis

All statistical analysis was done using GraphPad Prism. Unless indicated otherwise, data distributions were assumed to be normal, but this was not formally tested. Statistical comparison between indicated conditions was conducted using the two-sided Student’s t test or one-way ANOVA, as indicated in the figure legends. After running an ANOVA, the Tukey test was used to compare the mean with every other mean. Differences were considered significant if the p-value was <0.05.

## ACKNOWLEDGEMENTS

We thank Stephen P. Bell and Masayuki Onishi for comments on the manuscript, and members of the Lew Lab for stimulating discussions. This work was funded by NIH/NIGMS grant R35GM122488 to D.J.L.

## AUTHOR CONTRIBUTIONS

Conceptualization, review, and editing manuscript— A.C.E. Wirshing, and D.J. Lew. Data curation, investigation, methodology, visualization, validation, and formal analysis—A.C.E. Wirshing and M. Yan. Drafting of manuscript— A.C.E. Wirshing. Project administration, supervision, and resources— D.J. Lew

## ABBREVIATIONS

CV: Coefficient of Variation
ER: Endoplasmic reticulum
ER-PM: Endoplasmic reticulum-plasma membrane
GFP: Green fluorescent protein:
mNG: mNeonGreen
PM: Plasma membrane

## SUPPLEMENTAL FIGURE CAPTIONS

**Supplemental Figure 1:**
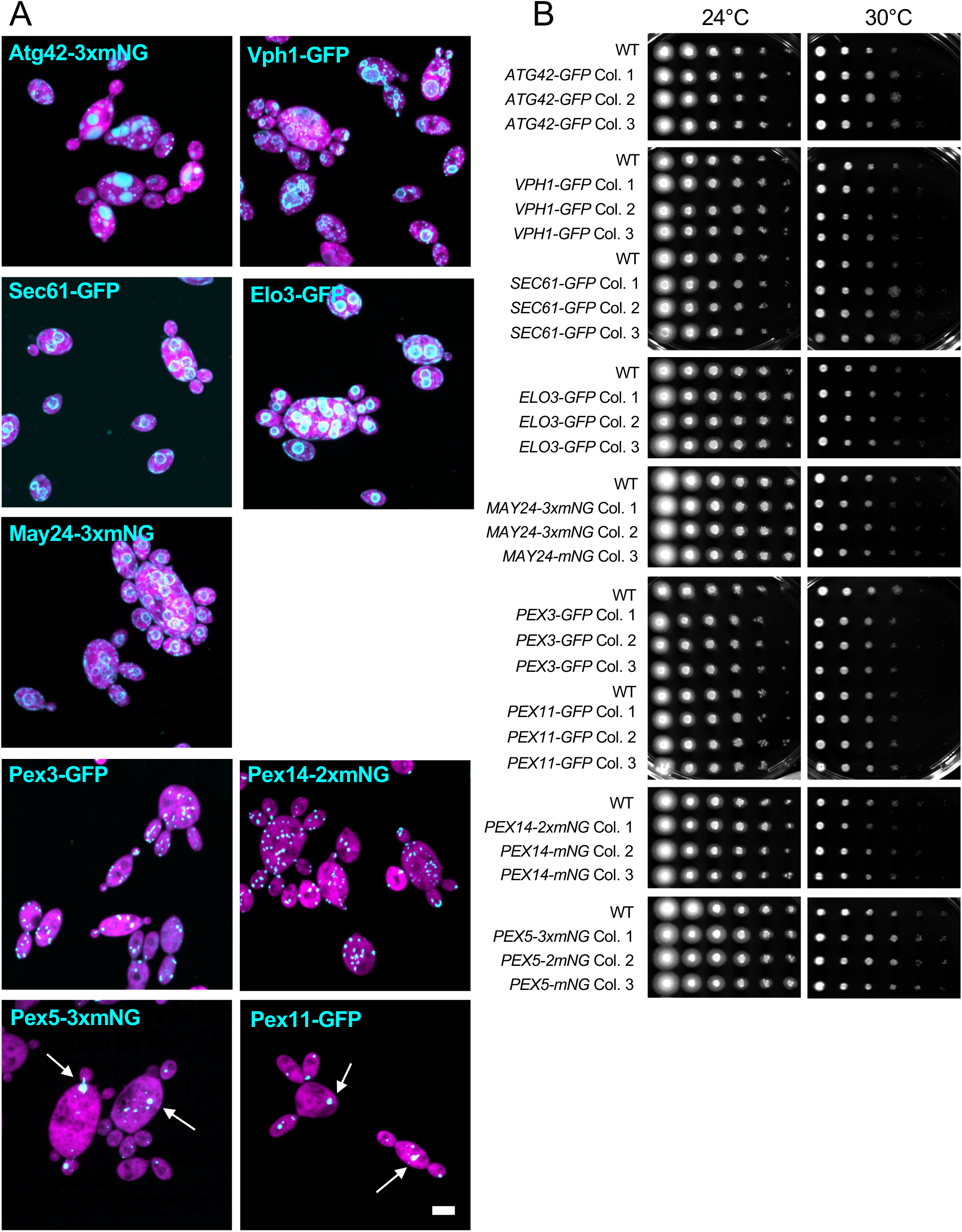
Characteriztion of organelle markers in *A. pullulans*. (**A**) Maximum intensity projections of confocal Z-series showing *A. pullulans* cells expressing a cytosol marker (3xmCherry, magenta) and each indicated organelle marker (cyan) tagged with GFP or mNG. The vacuole lumen was visualized using Atg42-3xmNG (DLY25963) and the vacuole membrane was visualized using Vph1-GFP (DLY24947). The ER was visualized using Sec61-GFP (DLY24944), Elo3-GFP (DLY27372), and May24-3xmNG (DLY27375). Peroxisomes were visualized using Pex3-GFP (DLY25001), Pex14-2xmNG (DLY27366), Pex11-GFP (DLY25004), and Pex5-3xmNG (DLY27370). Expression of Pex5-mNG or Pex11-GFP resulted in large clumps of peroxisomes (white arrows). Scale bar, 5 µm. (**B**) Growth assays on YPD showing 10-fold serial dilutions of three separate transformants with the indicated genotypes. Plates were grown for 48 h at the indicated temperatures.

**Supplemental Figure 2:**
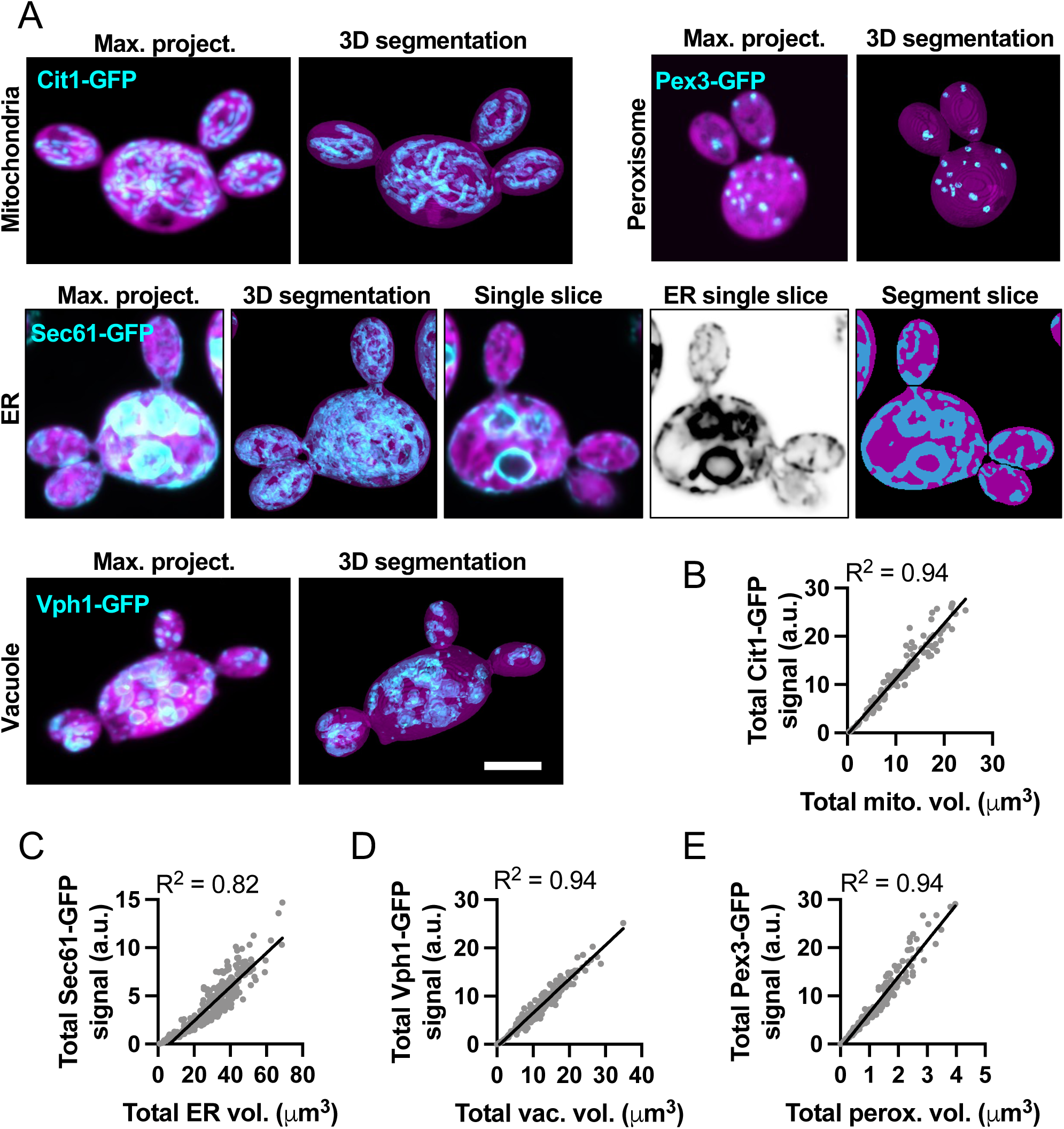
Quantification of organelle content. (**A**) Maximum intensity projections of confocal Z-series showing *A. pullulans* cells expressing a cytosol marker (3xmCherry, magenta) and different organelle markers in cyan: Cit1-GFP (DLY25312), Sec61-GFP (DLY24944), Vph1-GFP (DLY24947), and Pex3-GFP (DLY25001), labeling the mitochondria, ER, vacuole, and peroxisomes, respectively. To the right of each image we show the 3D segmentation used to measure cell and organelle volume. For the ER we also include a cross section to highlight cortical and internal ER structures. Scale bar, 5 µm. (**B-E**) Correlation between total organelle volume and total GFP signal measured in each bud for cells expressing different organelle markers: (**B**) mitochondria, (**C**) ER, (**D**) vacuole, and (**E**) peroxisome. R^2^ values are shown (n = 154, 304, 225, and 249 buds, respectively).

## SUPPLEMENTAL VIDEO CAPTIONS

**Supplemental video 1: Peroxisomes dynamically move between mother and bud.** Single confocal plane of a cell expressing the peroxisome marker Pex3-GFP (cyan) and cytoplasmic marker 3xmCherry (magenta). Images were acquired every 200 ms. Scale bar, 5 µm.

**Supplemental video 2: Organelle inheritance in multi-budded cells.** Maximum intensity projection of confocal time series of cells expressing a cytosol marker (3xmCherry, magenta) and different organelle markers in cyan: mitochondria (Cit1-GFP, DLY25312), ER (Sec61-GFP DLY24944), vacuole (Vph1-GFP DLY24947), and peroxisomes (Pex3-GFP DLY25001). Images were acquired every 5 min. Scale bar, 5 µm.

**Supplemental video 3: Some cells do not inherit peroxisomes.** Maximum intensity projection of confocal time series of cells expressing a cytosol marker (3xmCherry, magenta) and a peroxisome marker (Pex3-GFP, cyan) (DLY25001). On the left, peroxisomes enter the bud, and then return to the mother cell before cytokinesis (15 min) leaving the bud without peroxisomes. On the right, all peroxisomes are in the first bud at the time of cytokinesis (30 min). A second bud begins to emerge while there are still no peroxisomes in the mother cell (65 min). A dim Pex3-GFP puncta, presumptive newly forming peroxisome, appears in the mother cell (80 min). Newly formed peroxisomes stay in the mother cell and the second bud has none at the time of cytokinesis (125 min). Images were acquired every 5 min. Scale bar, 5 µm.

**Supplemental video 4: 3-color imaging of mitochondria, ER, and vacuole.** Individual slices of a confocal Z-series of wild-type cell stained with a mitochondria stain (magenta, MitoTracker) and expressing an ER marker (yellow, Sec61-3xmCherry) and vacuole marker (cyan, Vph1-3xmNG) (DLY26775). This accompanies data in Figure 7 in the main text. Scale bar, 5 µm.

**Supplemental Table 1.**

| Strain Name | Relevant Genotype |
| --- | --- |
| DLY24944 | *URA3:3xmCherry; SEC61-GFP:HYG<sup>R</sup>* |
| DLY24947 | *URA3:3xmCherry; VPH1-GFP:HYG<sup>R</sup>* |
| DLY25001 | *URA3:3xmCherry; PEX3-GFP:HYG<sup>R</sup>* |
| DLY25004 | *URA3:3xmCherry; PEX11-GFP:HYG<sup>R</sup>* |
| DLY25312 | *URA3:3xmCherry; CIT1-GFP:NAT<sup>R</sup>* |
| DLY25963 | *URA3:3xmCherry; ATG42-3xmNG:HYG<sup>R</sup>* |
| DLY26335 | *URA3:3xmCherry; VPH1-GFP:HYG<sup>R</sup>; pak1ΔNAT<sup>R</sup>* |
| DLY26338 | *URA3:3xmCherry; PEX3-GFP:HYG<sup>R</sup>; pak1ΔNAT<sup>R</sup>* |
| DLY26341 | *URA3:3xmCherry; SEC61-GFP:HYG<sup>R</sup>; pak1ΔNAT<sup>R</sup>* |
| DLY26344 | *URA3:3xmCherry; CIT1-GFP:NAT<sup>R</sup>; pak1ΔHYG<sup>R</sup>* |
| DLY26775 | *SEC61-3xmCherry:HYG<sup>R</sup>; VPH1-3xmNG:NAT<sup>R</sup>* |
| DLY27366 | *URA3:3xmCherry; PEX14-2xmNG:NAT<sup>R</sup>* |
| DLY27370 | *URA3:3xmCherry; PEX5-3xmNG:NAT<sup>R</sup>* |
| DLY27372 | *URA3:3xmCherry; ELO3-GFP:NAT<sup>R</sup>* |
| DLY27375 | *URA3:3xmCherry; MAY24-3xmNG:NAT<sup>R</sup>* |

